# Reassessing the Evidence for Universal School-age Bacillus Calmette Guerin (BCG) Vaccination in England and Wales

**DOI:** 10.1101/624916

**Authors:** Sam Abbott, Hannah Christensen, Ellen Brooks-Pollock

## Abstract

**Background:** In 2005, England and Wales switched from universal BCG vaccination against tuberculosis (TB) disease for school-age children to targeted vaccination of neonates. We assessed the quantitative evidence that informed this policy change.

**Methods:** We recreated a previous approach for estimating the impact of ending the BCG schools’ scheme in England and Wales, updating the model with parameter uncertainty. We investigated scenarios considered by the UK’s Joint Committee on Vaccination and Immunisation, and explored new approaches using notification data. We estimated the number of vaccines needed to prevent a single notification, and the average annual additional notifications caused by ending the BCG schools’ scheme.

**Results:** We found a 1.9% annual decrease in TB incidence rates best matched notification data. We estimate that 1600 (2.5-97.5% Quantiles (Q): 1300 - 2100) vaccines would have been required to prevent a single notification in 2004. If the scheme had ended in 2001, 302 (2.5-97.5% Q: 238 - 369) additional annual notifications would have occurred compared to if the scheme had continued. If the scheme ended in 2016, 120 (2.5-97.5% Q: 88 - 155) additional annual notifications would have occurred.

**Conclusions:** Our estimates of the impact of ending the BCG schools’ scheme were highly sensitive to the annual decrease in incidence rates. The impact of ending the BCG schools’ scheme was found to be greater than previously thought when parameter values were updated and notification data were used. Our results highlight the importance of including uncertainty when forecasting the impact of changes in vaccination policy.

**What is already known on this subject:**

- Targeted Bacillus Calmette Guerin (BCG) vaccination against TB is recommended in low incidence countries over universal vaccination.
- The impact of replacing universal BCG vaccination in England and Wales with a targeted programme in 2005 was assessed under the assumption of static declines in TB rates.
- The BCG Guerin vaccine was shown to be effective in the UK born in England, regardless of the age at which it was given. School-age vaccination maybe more beneficial in this population than in other settings.

**What this study adds:**

- Using notification data from England and Wales from 1973 to 2015, we estimate that the ending the BCG schools’ scheme likely resulted in more additional cases than was predicted.
- The inclusion of parameter uncertainty, and measurement error, allowed the uncertainty in the final estimates to be presented. Previously published estimates may have been spuriously precise.
- This study highlights the need for a more detailed evaluation of the 2005 change in BCG policy. In particular, the impact of including the introduction of targeted neonatal vaccination and capturing the long term, indirect, effects needs further study.

## Introduction

The Bacillus Calmette–Guérin (BCG) vaccine remains the only licensed vaccine for use against Tuberculosis (TB), although its use globally is controversial due to evidence of variable effectiveness,[1] and waning protection 10-15 years after vaccination.[2] Global usage of the BCG varies between no vaccination, universal vaccination, and high-risk group vaccination and may target either neonates or school-aged children.[3,4] The World Health Organization (WHO) recommends vaccination for all neonates as early as possible after birth in high burden settings, with vaccination in low burden settings being dependent on the country specific epidemiology of TB.[5] This recommendation is based on the strong evidence that the BCG is highly protective in children,[6,7] whilst its effectiveness has been shown to vary with latitude when given later in life.[8]

In England and Wales, universal school-aged vaccination was introduced after a MRC trial in the 1950s estimated BCG’s effectiveness at 80% in the ethnic White UK born population.[9] The policy remained in place until 2005, when England and Wales changed to targeted vaccination of neonates. The 2005 change in BCG vaccination policy was motivated by evidence of decreased transmission of TB, an increasing proportion of TB cases occurring in the non-UK born,[10] and modelling evidence that suggested stopping the BCG schools’ scheme would have minimal long term effects on incidence rates.[11] Due to the complex nature of both TB and the BCG vaccine, the ongoing impact of this change in policy is hard to directly estimate, with decision makers relying on expert opinion, evidence from surveillance data, and insight from modelling studies.

In 1987, an assessment of the school-age vaccination program was carried out in England and Wales.[11] This study was used, combined with a sensitivity analysis of notification rates, as supporting evidence by the Joint Committee on Vaccination and Immunisation (JVCI) BCG subgroup for the change in vaccination policy.[12,13] This papers aims to re-evaluate this modelling, and re-estimate the predicted impact of stopping the schools’ scheme. Whilst these results are retrospective they may be used by policy makers to assess some of the ongoing impact of ending the BCG schools’ scheme. In addition they highlight the importance of including uncertainty when forecasting the impact of changes in vaccination policy.

## Methods

### Modelling the impact of ending the BCG schools’ scheme

We implemented Sutherland et al.’s model for estimating the impact of ending the BCG schools’ scheme.[11] This model was based on data from TB notification surveys conducted in 1973, 1978, and 1983.[14] These were used to estimate incidence rates, stratified by BCG vaccination status, in the ethnic White UK born population of England and Wales aged 15-19 years old, 20-24 years old and 25-29 years old. Future incidence rates were forecast by assuming an annual decrease in incidence rates, which was based on historic trends (see supplementary information).[11,15] Primary impacts from ending the schools’ scheme were estimated by calculating the difference in incidence rates between the vaccinated and unvaccinated populations. Additional notifications from TB transmission were then calculated using a transmission chain model. This model was defined using the following steps,

1. Estimate the total expected number of secondary notifications (*T*) arising from any single primary notification using the following,

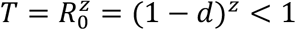 Where *R*_0_ is the expected number of secondary cases produced by a single infection in a completely susceptible population,[16] *d* is the percentage decay in notification rates, and *z* is the average interval between the notification of any individual and the notification of the patient who infected them.
2. Relate *T* to the number of notifications in each generation using the initial generation size (*x*) with the following power series,

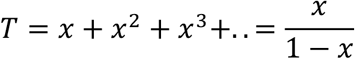
3. Estimate the expected average interval between each primary notification and all secondary notifications (*Z*). This is defined to be the sum of time to all notifications, weighted by the fraction in each generation, divided by the sum of all notifications. Mathematically this is,

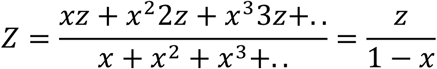 Implementing the model required several assumptions not detailed in [11]. Firstly, as incidence rates for those ineligible for the BCG schools’ scheme are not published, we assumed that they were equal to those in the unvaccinated population. In addition, in order to reproduce the distribution of cases due to transmission over time we introduced an additional parameter; the percentage of secondary cases due to a primary case in the first year after activation (*f*). The distribution of secondary cases (*N*) was then modelled as follows,

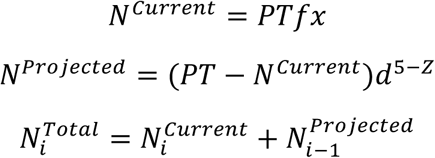

where *P* is the number of primary notifications, and *i* is the year of interest. We fitted this parameter using least squares to the original estimates of the total notifications due to ending the scheme under several scenarios, for several years. We validated our fitted model by comparing our results with those from the original implementation. Using the mean absolute percentage error, normalised by the orginal estimate.

### Updating model parameter estimates

Incidence rates were included as point estimates in [11]; in our updated model we included uncertainty in these rates. We estimated notifications for 1973, 1978, and 1983, using published incidence rates and population estimates. Samples were then generated using a Poisson distribution.[11,14] These samples were then used to estimate a distribution of incidence rates. Sutherland et al. assumed a serial interval of 2 years between linked infections. We updated this assumption using an estimate from a recent study in the Netherlands of 1.44 (95% CI 1.29 to 1.63) years.[17]

We considered the original assumption of a 9% annual decrease in incidence rates as well as three scenarios based on those considered by the JVCI BCG subgroup:[12,13] these were a 3.9% decrease, a 1.9% decrease, and no change annual in incidence rates. We also estimated the annual decrease in incidence rates in the ethnic White UK born using two proxy measures. The first proxy measure was the annual change in notifications in England and Wales, which was estimated using data from Public Health England (PHE). The standard deviation (SD) of this measure was then calculated using the prop.test function in R.[18] The second proxy used was the annual decrease in the UK born age-specific incidence rates in the English population. These were calculated using notification data from the Enhanced Tuberculosis surveillance system (ETS) and the June Labour Force Survey.[10] Incidence rates (with SDs) were estimated using the epiR package.[19]. Uncertainty was incorporated by sampling from a normal distribution for both proxy measures. Data collection for the ETS began in 2000, we therefore estimated incidence rates between 1984 and 1999, and for the years between notifications surveys (1974-1977 and 1979-1892), using local regression for each sample fitted to the incidence rates published in [11], and our estimated incidence rates from 2000 on-wards. For years prior to 1973 the annual decreases were assumed to be the mean of the previous 3 years of data. For both proxy measures the annual decreases in incidence rates post 2016 were assumed to be the average of the estimates in 2013, 2014, and 2015.

### Statistical analysis

For each scenario, we ran the model until 2028 with 10,000 parameter samples. We tested the difference between scenarios using the Mann-Whitney test for both the number of vaccines needed to prevent a single case in 15 years after vaccination for a cohort aged 13- 14 years old at vaccination and the total number of additional notifications caused by ending the BCG schools’ scheme. As in [11] a 15 year time horizon was used with 5 year intervals. The year closest to the year of the change in vaccination policy (2005), which had model estimates, was used as the baseline.

## Results

### Model validation

Our model produced results that were comparable with those from [11] (supplementary table S1). When estimating the total notifications from ending the BCG schools’ scheme at different times in ethnic White UK born adults aged 15-29 years old in England and Wales our model had a median absolute error of 2.03% (2.5-97.5% Q: 0.00% - 3.72%) and a maximum absolute error of 3.91% when compared to [11].

### Annual change in TB incidence rates

We found that the assumption of a 9% annual decrease in incidence rates in the ethnic White UK born was not comparable to estimates using either notification data or age-specific incidence rates in the time period studied (supplementary figure S1). The median annual decrease estimated using notifications was 3.13% (2.5-97.5% Quantiles (Q): −8.32% - 11.45%), with a maximum of 15.13% (2.5-97.5% Q: 14.23% - 16.04%) in 1987 and a minimum of −10.18% (2.5-97.5% Q: −10.82% - −9.52%) in 2005. Using age-specific incidence rates we estimated the median annual decrease in incidence rates for 15-19 year olds was 1.62% (2.5-97.5% Q: -40.38% - 39.89%), 3.15% (2.5-97.5% Q: −33.93% - 38.25%) for 20-24 year olds, and 2.66% (2.5-97.5% Q: −36.37% - 37.29%) for 25-29 year olds. There was substantial variation between years and a high degree of uncertainty.

### Vaccines required to prevent a single notification

We found that updating parameter values, and incorporating uncertainty, did not alter the number of vaccines required to prevent a single notification within 15 years in a cohort vaccinated at school-age, when the annual decrease in TB incidence rates was assumed to be 9% (supplementary table S2). However, the updated estimate had a wide range (15000 (2.5-97.5% Q: 12000 - 19000) vaccines required in 2004). As the assumed annual decrease in incidence rates was reduced the number of vaccines required to prevent a single notification also reduced; an estimated 1600 (2.5-97.5% Q: 1300 - 2100) vaccines were required to prevent a case in 2004 when the annual decrease was assumed to be 1.9%. Estimates of the number of vaccines required to prevent a notification were comparable but not equivalent when the annual decrease was estimated using notifications (1400 (2.5- 97.5% Q: 1100 - 1700), P: 0.077) and age-specific incidence rates (1500 (2.5-97.5% Q: 450 - 5000), P: 0.083). The estimate using incidence rates had a high degree of uncertainty (figure 1). The number of vaccines required increased slightly over time with 1800 (2.5- 97.5% Q: 1500 - 2300) required in 2009, 2000 (2.5-97.5% Q: 1600 - 2500) required in 2014, and 2200 (2.5-97.5% Q: 1800 - 2700) required in 2019 when an annual decrease of 1.9% in incidence rates was assumed.

**Figure 1:**
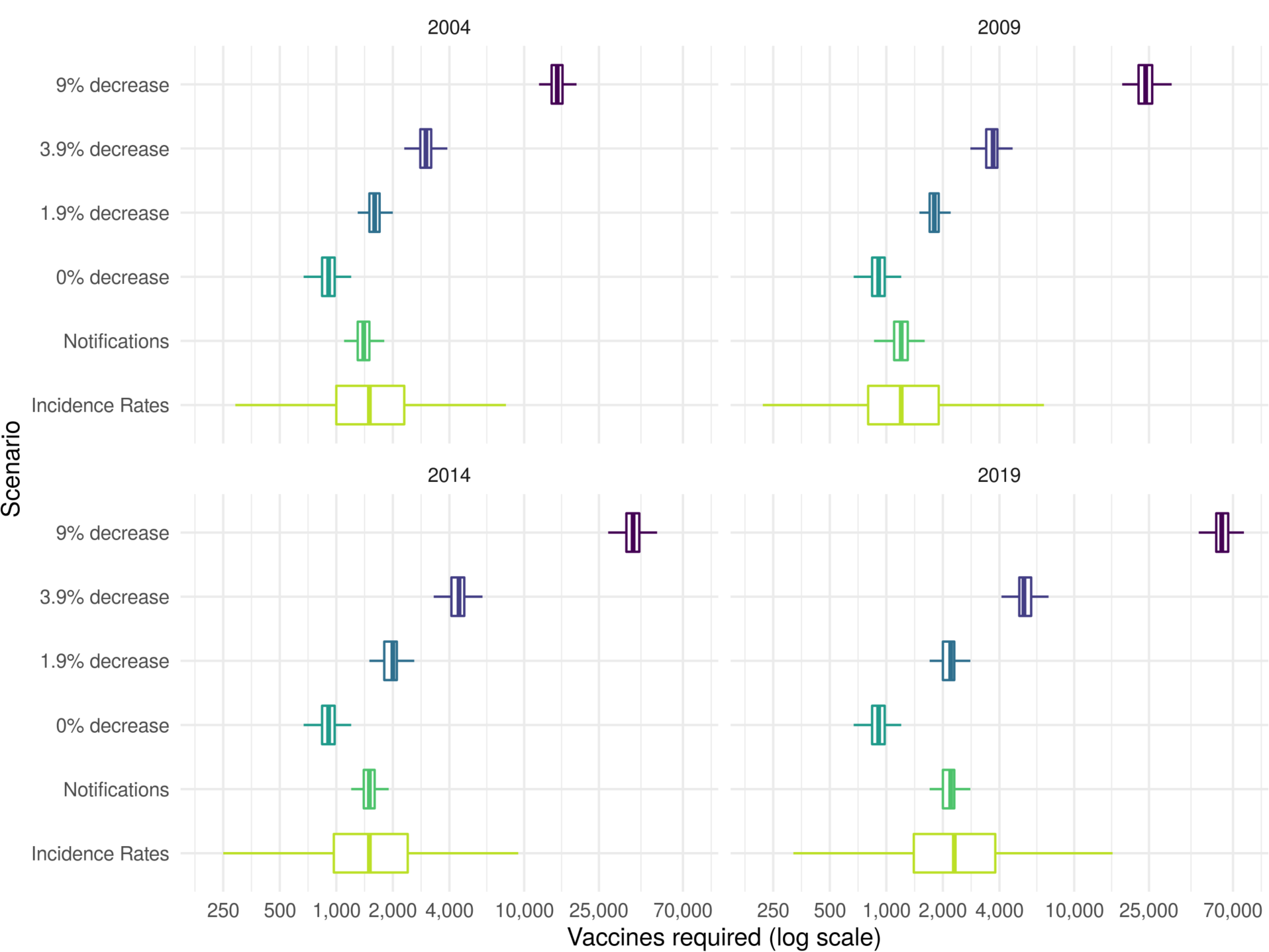
Vaccines required in a cohort of those vaccinated at school-age to prevent a single case of Tuberculosis within 15 years of vaccination in 2004, 2009, 2014, or 2019. The years presented were dictated by the 5 year timestep of the model. The percentage annual decrease scenarios considered were based on those considered by the JVCI BCG subgroup, with the addition of a scenario using aggregate notification data and a scenario using estimates of age-specific incidence rates in the UK born. Each boxplot summarises the output of 10,000 model simulations for each scenario. Outliers have been omitted for clarity.

**Figure 2:**
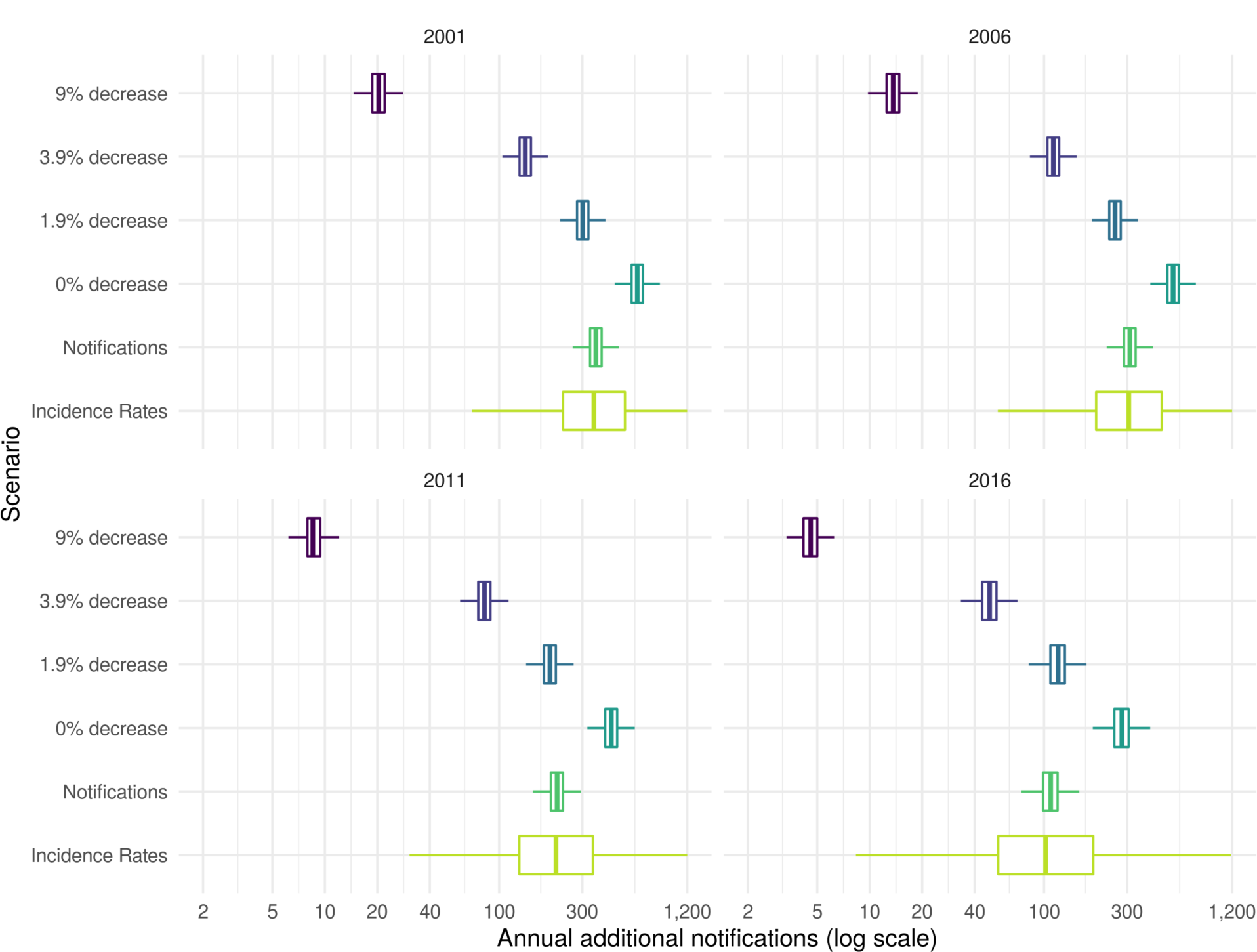
Annual additional notifications in 15-29 year olds from stopping the BCG schools’ scheme in 2001, 2006, 2011, and 2016 until 2028. The years presented were dictated by the 5 year timestep of the model. The percentage annual decrease scenarios considered were based on those considered by the JVCI BCG subgroup, with the addition of a scenario using aggregate notification data and a scenario using estimates of age-specific incidence rates in the UK born. Each boxplot summarises the output of 10,000 model simulations for each scenario. Outliers have been omitted for clarity.

### Average annual additional cases from ending the BCG schools’ scheme at various dates

We found that updating parameter values, and incorporating uncertainty, did not alter the average annual additional notifications from stopping the BCG schools’ scheme when the annual decrease was assumed to be 9% (supplementary table S3). There was a large degree of uncertainty in this estimate with 20 (2.5-97.5% Q: 16 - 25) notifications prevented annually if vaccination was stopped in 2001. As the assumed annual decrease in incidence rates was reduced the annual number of additional notifications prevented increased with 302 (2.5-97.5% Q: 238 - 369) notifications prevented annually when the annual decrease was assumed to be 1.9% and vaccination stopping in 2001. There was some evidence that the average annual number of notifications prevented was greater when the annual decrease was estimated using notifications (359 (2.5-97.5% Q: 282 - 439), P: 0.083) and age-specific incidence rates (359 (2.5-97.5% Q: 102 - 1332), P: 0.083), compared to an assumed annual decrease of 1.9% (figure 1). The estimate made using incidence rates again had a high degree of uncertainty. When an annual decrease of 1.9% was assumed the number of notifications prevented annually reduced with time: 255 (2.5- 97.5% Q: 201 - 313) from ending vaccination in 2006; 196 (2.5-97.5% Q: 152 - 242) from ending vaccination in 2011, and 120 (2.5-97.5% Q: 88 - 155) from ending vaccination in 2016.

## Discussion

The existing method for estimating the impact of the BCG schools’ scheme produced uncertain estimates of the impact of ending the scheme in all years evaluated when parameter uncertainty and measurement error were included. Updating the annual decrease in TB notifications based on both notifications and using age-specific incidence rates resulted in increased TB cases due to ending universal school-age vaccination in all years considered. This resulted in fewer vaccines required to prevent a single notification in those vaccinated and an increase in the number of annual additional notifications from ending the scheme. A scenario with a 1.9% annual decrease in incidence rates was most comparable to our results based on notifications.

This study reassesses a key piece of the quantitative evidence used to motivate the change in BCG vaccination policy in 2005. Our results provide new insights into the uncertainty of the previously published model predictions by including parameter uncertainty and measurement error. As the data on incidence rates in the ethnic White UK born in England and Wales were not available we considered two approaches to proxy them, and investigated multiple scenarios based on those explored by the JVCI BCG subgroup. The simulation approach used here is not the most accurate method for assessing the impact of ending the BCG schools’ scheme. However, it does provide an estimate that is based on the available data and on the framework used to inform policy making. This allowed the strength of some the quantitative evidence used in the decision-making process to be assessed. This would not have been possible if the impact had been assessed using only the observed data. A weakness of the modelling framework used in this study is that it did not include the whole population or age groups outside those directly affected by vaccination. Furthermore, heterogeneous mixing between these groups is likely to be important. The exclusion of these factors means that our results are conservative. A final limitation is that this study only considers the impact of ending the BCG schools’ scheme and not the impact of the introduction of the targeted neonatal vaccination program. This should be considered when evaluating the change in policy as a whole.

Little work has been done to assess the impact of the 2005 change in BCG vaccination policy or to assess the quantitative evidence used in decision making. However, multiple studies have evaluated the cost effectiveness of various BCG programs and the impact of switching between them. A cluster-randomised trial in Brazil found that BCG vaccination of those at school-age was cheaper than treatment and would prevent one TB case per 381 vaccinations even with a vaccine effectiveness of only 34% (8-53%).[20] This is substantially fewer than our estimate of 2000 (2.5-97.5% Q: 1600 - 2500) vaccines required to prevent a single notification within 15 years in 2014 (this was the most comparable year from our study). However, the same trial found that for regions close to the equator BCG effectiveness was low in school-age children but unchanged in neonates,[21] highlighting the importance of considering the BCG vaccines reduced effectiveness near the equator when determining vaccination policy.[22] There is also some research which supports universal re-vaccination of those at school-age, in countries with high incidence and universal vaccination of neonates, as it may be cost effective when BCG effectiveness is moderate to high.[21,23] There is some evidence that targeted vaccination of high risk neonates maybe more cost effective than universal vaccination of neonates.[24,25] However, a study in Sweden found that incidence rates in Swedish-born children increased slightly after universal vaccination of neonates was discontinued in favour of targeted vaccination.[26] In France, which switched from universal vaccination of neonates to targeted vaccination in 2007, it has also been shown that targeted vaccination reduced coverage in those most at risk.[27] Targeted vaccination may not be more cost effective that universal vaccination when possible reductions in transmission are considered. Our study indicated that a substantial number of cases due to transmission may be preventable if universal school-age BCG vaccination was still in place. This result is dependent on the effectiveness of BCG vaccination when given later in life, for which there is good evidence in the ethnic White UK born.[9] We did not consider neonatal vaccination which would be less impacted by BCG’s effectiveness reducing when given later in life, but would also be less likely to result in the same reductions in ongoing transmission.

This study indicates that some of the evidence used to justify the 2005 change in BCG vaccination policy may have underestimated the impact of ending the scheme. It highlights the importance of including both parameter and measurement error, as excluding these sources of variation may lead to spuriously precise results. In addition, our exploration of the assumptions used to estimate the annual change in TB incidence rates in the ethnic White UK born illustrates the structural impact of assuming an annual decrease in TB incidence rates. More realistic estimates of the annual decrease in incidence rates resulted in a greatly increased impact of ending the BCG schools’ scheme. Policy makers should consider these updated estimates when assessing the role of BCG vaccination in those at school-age.

This study has reassessed some of the evidence previously used in decision making, updating the approach with new data. However, as 15 years of detailed surveillance data have been collected since the ending of the BCG schools’ scheme it is now possible to use regression-based approaches to estimate the direct impact on incidence rates of ending the BCG schools’ scheme. These approaches could also be used to estimate the impact of vaccinating high-risk neonates, which may outweigh any negative impacts of ending the BCG schools’ scheme. In addition, the development, and use, of a transmission dynamic model would allow the more accurate estimation of indirect effects and the forecasting of long-term impacts.

## Acknowledgements

The authors thank the TB section at Public Health England (PHE) for maintaining the Enhanced Tuberculosis Surveillance (ETS) system; all the healthcare workers involved in data collection for the ETS.

## Contributors

SA, HC, and EBP conceived and designed the work. SA undertook the analysis with advice from all other authors. All authors contributed to the interpretation of the data. SA wrote the first draft of the paper and all authors contributed to subsequent drafts. All authors approve the work for publication and agree to be accountable for the work.

## Funding

SEA, HC, and EBP are funded by the National Institute for Health Research Health Protection Research Unit (NIHR HPRU) in Evaluation of Interventions at University of Bristol in partnership with Public Health England (PHE). The views expressed are those of the author(s) and not necessarily those of the NHS, the NIHR, the Department of Health or Public Health England.

## Conflicts of interest

HC reports receiving honoraria from Sanofi Pasteur, and consultancy fees from AstraZeneca, GSK and IMS Health, all paid to her employer.

## Accessibility of data and programming code

The code for the analysis contained in this paper can be found at: DOI:10.5281/zenodo.2583056

## Notes

https://github.com/seabbs/AssessBCGPolicyChange/

