## Supplementary Information for "Reassessing the Evidence for Universal School-age Bacillus Calmette Guerin (BCG) Vaccination in England and Wales"

### Recreating the estimation model of Sutherland et al.

#### Estimating notification rates

The effectiveness of the BCG vaccine was originally estimated by an MRC trial in 1953 at 80% in the United Kingdom [10]. As follow up to this trial members of the MRC bio-statistics unit conducted a series of notification surveys attempting to ascertain any change in effectiveness, as well as acting as an estimate of notification rate across different demographics.[15] Notification surveys were carried out at 5 year intervals in 1973, 1978 and 1983; looking at those aged 15-24 years old in England and Wales. For the 1983 survey records of BCG status, Tuberculin status and ethnicity were extracted from the records of notifying physicians and the records of the local health and education authorities. Totals, across the study period, were then aggregated for the following group; Tuberculin negative and BCG positive, Tuberculin negative and BCG negative, Tuberculin positive and not vaccinated and those who did not participate. These totals were then combined with the population estimates for each cohort at 13 years of age to estimate the ethnic make up of the population, and to construct notification rates for each category. Data drawn from a range of sources including: Office of National Statistics data; annual local authority returns for total tuberculin test results; BCG vaccinations in the schools' scheme; and the Labour force survey (1983),

For 1983 this corresponded with 874 notifications in the ethnic White UK born in England and Wales, between 15-24, with participation at 80%. As the number of Tuberculin negative subjects not given BCG was unreported this was estimated at 1.9% of all vaccinated with the BCG. Allowances were also made for those already vaccinated prior to the schools' scheme. See [15] for full details of the survey construction and the additional assumptions used to give similar estimates for both the 1973 and 1978 surveys. The findings of these surveys were as follows; in the ethnic White population notification rates had fallen by an annual average of 9% and BCG efficacy had remained high.[12,16]

Evidence suggests that the BCG vaccine has a high efficacy for at least the first 15 years after vaccination, therefore the authors extrapolated from the data on the 20-24 cohort to a theoretical 25-29 year old cohort. Data on the notifications for the first 6 months of the 1983 survey was available and this was then scaled using the ratio of notifications from this age group against the total number of notifications recorded in that year. Population estimates from the 20-24 cohort were adjusted for all causes of mortality (0.34%). Migration was ignored. The tuberculin positive cohort had a sharp decline in the previous two cohorts, therefore it was assumed that this continued. Lastly the efficacy was estimated as being that seen in the 20-24 cohort but with the same decline in protection seen between the last two cohorts. These assumptions allowed notification rates to be estimated for the 25-29 year old population, resulting in a complete cohort over the projected 15 years of BCG effectiveness.

### Construction of forward estimates

Based on these estimated notification rates Sutherland et al. then sought to quantify the ongoing risk of developing notified tuberculosis, projected forward in time, for both the vaccinated and unvaccinated populations. To construct these estimates several key assumptions, based on the results seen in the previous surveys, were made. Firstly, it was assumed that efficacy was not degrading within the ethnic White population and therefore historic estimates would continue to apply into the future. Additionally, it was assumed that the decay of 9% in notification rates, across all ethnic White populations, would continue indefinitely.

These assumptions allow the notification rates in both the BCG vaccinated and unvaccinated groups to be projected forward in time. By assuming that the schools' scheme is responsible for the observed variation between vaccinated and unvaccinated rates the rate of prevented cases can then be estimated. By scaling this against a cohort of 100,000 13 year olds the number of prevented cases over a 15 year period can be projected for each cohort. By dividing the total number in a given cohort by the number of prevented cases the estimated number of vaccines required to prevent a single case in the 15 year period can then be calculated.

To estimate the total number of prevented notifications, for each cohort, in England and Wales the total number receiving the BCG and the coverage of the schools' scheme was required. The coverage of the BCG schools' scheme was estimated from annual reports of the DHSS and was assumed to be 75% for all years. The number of BCG vaccines given each year was estimated from the DHSS returns for the years 1967 to 1981, it was then taken as 75% of the estimated ethnic White population aged 13 years from 1982-1996, for each 5 year period thereafter it was assumed to be 2.1 million.

Using the above information and the projected differences between vaccinated and unvaccinated notifications allows the number of prevented notifications for each age group to be found for each year. This can then be combined to give the total number of prevented notifications for those aged between 15-29. To properly understand these estimates, estimates of the projected yearly notifications if the scheme continues are required. These totals were derived from the vaccinated and unvaccinated rates supplemented with similar projections from the tuberculin positive or otherwise ineligible sourced from the 1983 BCG survey.[15]

### Model validation

The percentage of cases in the first year was fitted to the Sutherland et al. estimates using the least squares method, such that $f = 0.764$.

*Supplementary Table S1: Comparison of results published by Sutherland et al. vs. our recreated model. This table shows the total notifications including primary and secondary effects from ending the BCG schools scheme at various times in white adults aged 15-29 years old in England and Wales.*

| Year of Ending Scheme | | | 1988 | | | 1993 | | | 1998 | | | 2003 | | | 2008 | | | 2013 |
|---|---|---|---|---|---|---|---|---|---|---|---|---|---|---|---|---|---|---|
| | Original | Recreated | Difference | Original | Recreated | Difference | Original | Recreated | Difference | Original | Recreated | Difference | Original | Recreated | Difference | Original | Recreated | Difference |
| 1986 | 288 | 296 | 8 | 226 | 226 | 0 | 208 | 205 | -3 | 181 | 175 | -6 | 128 | 123 | -5 | 80 | 78 | -2 |
| 1991 | 288 | 296 | 8 | 165 | 166 | 1 | 130 | 131 | 1 | 128 | 126 | -2 | 115 | 111 | -4 | 80 | 78 | -2 |
| 1996 | 288 | 296 | 8 | 165 | 166 | 1 | 90 | 91 | 1 | 77 | 76 | -1 | 80 | 80 | 0 | 72 | 70 | -2 |

Results

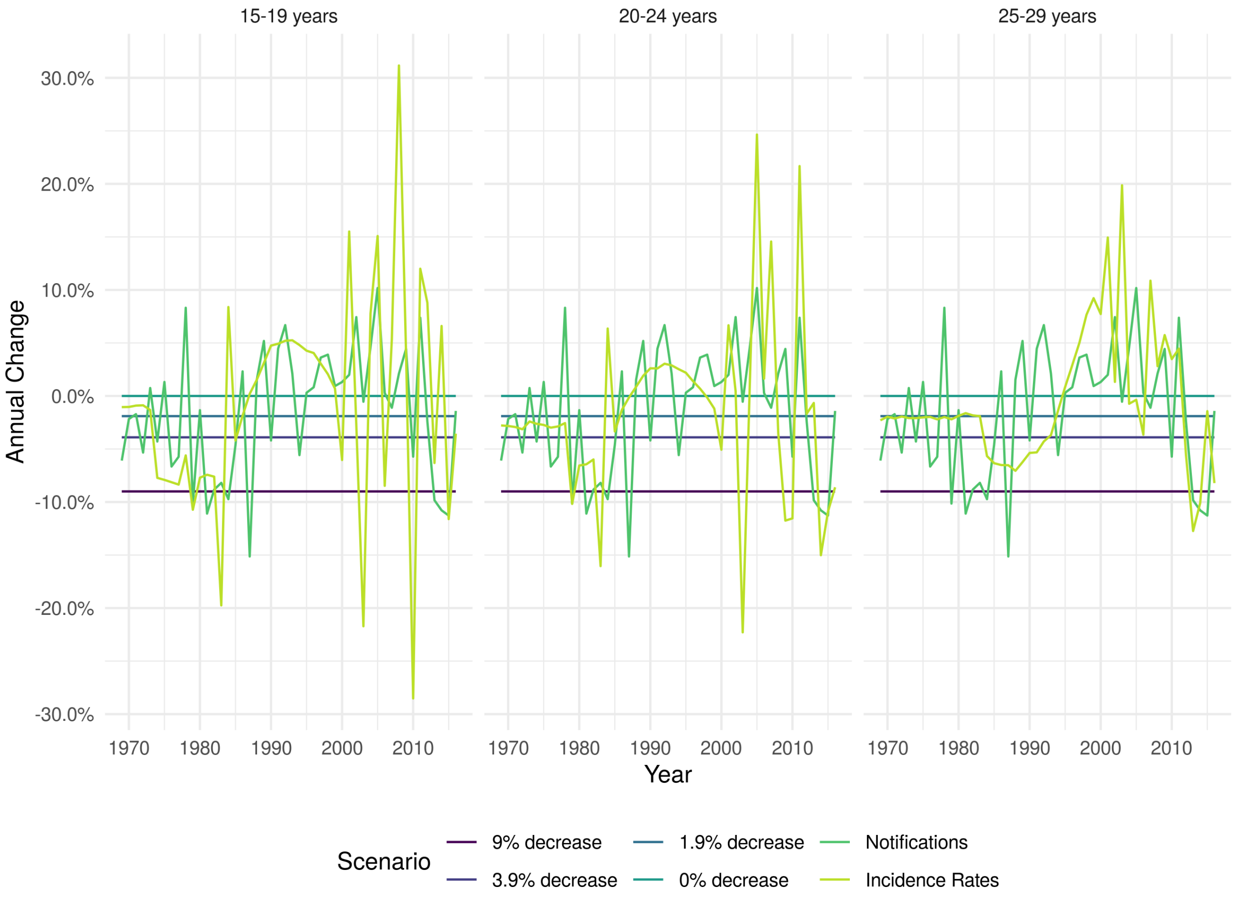


*Supplementary Figure S1: Annual percentage change in ethnic White UK born incidence rates for those aged 15-19, 20-24, and 25-29 years old under different scenarios. For the notification and incidence rate scenarios each line represents the median of 10,000 parameter samples.*

*Supplementary Table S2: The median number (with the 2.5% and 97.% quantiles) of vaccines required to prevent a single case of Tuberculosis within 15 years in a ethnic White UK born adult vaccinated at 13 years old.*

| Year of Vaccination | 9% decrease (original parameters) | 9% decrease | 3.9% decrease | 1.9% decrease | 0% decrease | Notifications | Incidence Rates |
|---|---|---|---|---|---|---|---|
| 1969 | 460 | 460 (390 - 550) | 460 (390 - 550) | 460 (390 - 550) | 460 (390 - 550) | 460 (390 - 550) | 460 (390 - 550) |
| 1974 | 940 | 940 (780 - 1200) | 880 (730 - 1100) | 860 (710 - 1100) | 830 (680 - 1000) | 860 (710 - 1100) | 870 (640 - 1100) |
| 1979 | 1400 | 1400 (1100 - 1800) | 1100 (910 - 1400) | 1000 (820 - 1300) | 910 (740 - 1200) | 1000 (860 - 1300) | 1100 (660 - 1700) |
| 1984 | 2200 | 2300 (1800 - 2900) | 1400 (1100 - 1700) | 1100 (900 - 1400) | 910 (740 - 1200) | 1600 (1300 - 2000) | 1500 (730 - 2900) |
| 1989 | 3600 | 3600 (2900 - 4600) | 1700 (1300 - 2100) | 1200 (990 - 1500) | 910 (740 - 1200) | 1800 (1500 - 2300) | 1700 (760 - 4000) |
| 1994 | 5800 | 5800 (4700 - 7400) | 2000 (1600 - 2600) | 1300 (1100 - 1700) | 910 (740 - 1200) | 1700 (1400 - 2200) | 1600 (630 - 4200) |
| 1999 | 9300 | 9300 (7600 - 12000) | 2500 (2000 - 3100) | 1500 (1200 - 1900) | 910 (740 - 1200) | 1600 (1300 - 2000) | 1500 (510 - 4200) |
| 2004 | 15000 | 15000 (12000 - 19000) | 3000 (2500 - 3800) | 1600 (1300 - 2100) | 910 (740 - 1200) | 1400 (1100 - 1700) | 1500 (450 - 5000) |
| 2009 | 24000 | 24000 (19000 - 30000) | 3700 (3000 - 4600) | 1800 (1500 - 2300) | 910 (740 - 1200) | 1200 (960 - 1500) | 1200 (330 - 4300) |
| 2014 | 38000 | 38000 (31000 - 49000) | 4500 (3600 - 5700) | 2000 (1600 - 2500) | 910 (740 - 1200) | 1500 (1200 - 1900) | 1500 (370 - 6000) |
| 2019 | 61000 | 61000 (50000 - 78000) | 5400 (4500 - 6900) | 2200 (1800 - 2700) | 910 (740 - 1200) | 2200 (1800 - 2700) | 2300 (470 - 10000) |
| 2024 | 98000 | 98000 (80000 - 130000) | 6600 (5400 - 8400) | 2400 (1900 - 3000) | 910 (740 - 1200) | 3200 (2600 - 4100) | 3300 (530 - 18000) |

*Supplementary Table S3: The median number (with the 2.5% and 97.% quantiles) of additional annual notifications due to ending the BCG schools' scheme in selected years.*

| Year Ending Scheme | 9% decrease (original parameters) | 9% decrease | 3.9% decrease | 1.9% decrease | 0% decrease | Notifications | Incidence Rates |
|---|---|---|---|---|---|---|---|
| 1971 | 204 | 203 (162 - 247) | 391 (310 - 476) | 557 (440 - 678) | 827 (652 - 1009) | 531 (421 - 646) | 548 (273 - 1254) |
| 1976 | 158 | 157 (123 - 193) | 350 (276 - 428) | 525 (414 - 643) | 816 (642 - 998) | 497 (391 - 607) | 520 (236 - 1282) |
| 1981 | 97 | 97 (76 - 119) | 282 (223 - 346) | 462 (364 - 565) | 767 (604 - 938) | 434 (341 - 531) | 462 (181 - 1275) |
| 1986 | 61 | 61 (48 - 75) | 231 (182 - 283) | 410 (323 - 501) | 724 (570 - 886) | 406 (319 - 496) | 427 (151 - 1282) |
| 1991 | 43 | 43 (34 - 53) | 200 (158 - 245) | 378 (298 - 462) | 703 (554 - 859) | 399 (313 - 487) | 419 (138 - 1336) |
| 1996 | 30 | 30 (23 - 36) | 170 (134 - 208) | 341 (269 - 418) | 668 (526 - 817) | 384 (301 - 469) | 397 (122 - 1354) |
| 2001 | 20 | 20 (16 - 25) | 141 (111 - 173) | 302 (238 - 369) | 620 (489 - 758) | 359 (282 - 439) | 359 (102 - 1332) |
| 2006 | 14 | 14 (11 - 17) | 113 (89 - 139) | 255 (201 - 313) | 550 (433 - 673) | 311 (244 - 380) | 314 (84 - 1265) |
| 2011 | 9 | 9 (6 - 11) | 82 (64 - 102) | 196 (152 - 242) | 440 (343 - 542) | 215 (167 - 265) | 215 (51 - 1026) |
| 2016 | 5 | 5 (3 - 5) | 49 (35 - 63) | 120 (88 - 155) | 280 (205 - 358) | 109 (80 - 140) | 103 (15 - 755) |
| 2021 | 2 | 2 (1 - 3) | 20 (13 - 29) | 52 (33 - 74) | 126 (78 - 176) | 39 (24 - 54) | 41 (3 - 491) |
